# First indication of acetylcholine-based communication in honeybee haemocytes and its modulation by a neonicotinoid insecticide

**DOI:** 10.1101/105700

**Authors:** T. Pamminger, K. Basley, D Goulson, WOH Hughes

## Abstract

There is growing concern that some managed and wild insect pollinator populations are in decline, potentially threatening biodiversity and sustainable food production on a global scale. In recent years, there has been increasing evidence that sub-lethal exposure to neurotoxic, neonicotinoid pesticides can negatively affect pollinator immunocompetence and amplify the effects of diseases, likely contributing to pollinator declines. Here we show that a range of non-neural tissues and haemocytes of the honeybee *Apis mellifera* express the nicotinic acetylcholine receptor that is the target of neonicotinoids. In addition, we demonstrate that the haemocytes, which form the cellular arm of the innate immune system, actively synthesize acetylcholine. This suggests the presence of a neural-independent acetylcholine-based immune signalling system in insects similar to that found in vertebrates. Lastly we establish that field-relevant doses of the neonicotinoid insecticide clothianidin alter this communication system. These findings provide a novel, mechanistically informed framework to understand the numerous siede-effects on insects of sub-lethal pesticide exposure, including immunosuppression. They support the growing evidence for acetylcholine-based immune regulation in invertebrates that operates independently of the nervous system.

## Introduction

In an attempt to meet the ever-increasing demand for pollination services, globalized pollinator-trade has led to the spread of pollinator diseases around the world, threatening managed and wild pollinator populations^1–3^. While healthy pollinator communities are sometimes able to contain such emerging diseases, additional stressors can compromise pollinator immunity, causing lethal epidemics^4–6^. One prominent factor directly impairing pollinator immunity is their exposure to sub-lethal doses of neurotoxic pesticides such as neonicotinoids^7–9^. While the demonstrated detrimental effects of neurotoxic pesticides on pollinator behaviour ^10,11^ and navigation ^12,13^ are intuitive, the strong immunosuppressive effects of these neurotoxic pesticides remain unexplained^14^.

In vertebrates it is well established that the immune system has a close regulatory connection with the nervous system^15^. In particular, the ancient cholinergic signalling system based on acetylcholine (*ACh*) has been demonstrated to perform a pivotal role in maintaining homeostasis of the immune system^16,17^. In recent years, evidence for a functionally similar *ACh*-based immune regulatory network has emerged in a handful of invertebrate model systems^18–20^. In particular, haemocytes, the cellular arm of the invertebrate immune system, have been demonstrated to not only express subunits of the muscarinic (*mAChR*)^21^ and nicotinic acetylcholine receptors (*nAChR*)^22^, but also to directly respond to the presence of *ACh*^23^. Since neonicotinoids target *nAChR* receptors with high affinity^24^ the presence of a neural-independent, *ACh* based communication system in the innate immune system of pollinators could provide a direct mechanistic link between neonicotinoids and immunosuppression.

In this study we investigate if non-neural immune-relevant tissues (fatbody, midgut and haemocytes) of the honeybee *Apis mellifera*: 1) express *nAChR* subunits, 2) synthesize *ACh*, and 3) respond to field-realistic doses of neonicotinoids. If confirmed, this would provide a mechanistic framework to directly explain the hitherto puzzling immunosuppressive effects of sub-lethal pesticide exposure observed in pollinators.

## Results

We find that all tissues investigated express a different subset of *nAChR* subunits (Pseudo-F_3,23_ = 7.76, P < 0.001, Fig. 1A-D, Fig 2). A pairwise comparison indicates that all four tissues (fat body, haemocyte, midgut, brain) express a unique blend of subunits (all comparisons t > 1.92, P < 0.006; for details see Electronic Supplementary Material (ESM) Table S1, Fig. 1A-D, Fig. 2). A PERMANOVA using Euclidian distance yielded very similar results (see Table S1). The SIMPER analysis indicates that subunit α7 is the most highly expressed subunits in brain tissue separating it from all other tissues (Table S2, Fig. 1D). In contrast subunit α9 and β2 are mostly expressed in the fatbody making it different from the other tissues (Table S2, Fig. 1A). Haemocytes, similarly to brain tissue, exhibit biased expression of α2 and α7 (Fig. 1B). Midgut tissue exhibits low expression levels of all subunits (Fig. 1C). The tissues also differed in terms of *ACh* expression measured as choline acetyltransferase (*ChAT*) expression (X^2^ = 21.96. P < 0.001, Fig. 2A), with brain tissues having the highest expression compared to all other tissues (all P < 0.001, Fig. 2A) followed by haemocytes (all P < 0.001, Fig. 2A), with very little expression activity in the midgut and fatbody (P > 0.05, Fig. 2A). Over the 24 h experiment, bees in the clothianidin treatment group consumed on average 0.44 ± 0.16 ng of clothianidin. Treatment groups did not differ in terms of survival (z = -0.26, P = 0.79). Control and pesticide-exposed bees consumed a similar amount of sucrose solution (W = 114, P = 0.95), and we were able to extract similar amounts of haemolymph from both treatment groups (W = 23, P = 0.82). When looking at the amount of total RNA extracted from bee haemolymph we find no difference between treatment groups (totalRNA/µl haemolymph W = 12, P = 0.23). However, we found that *ChAT* expression was significantly increased in haemocytes of bees exposed to clothianidin (W = 38, P = 0.014, Fig. 2B).

**Figure 1:**
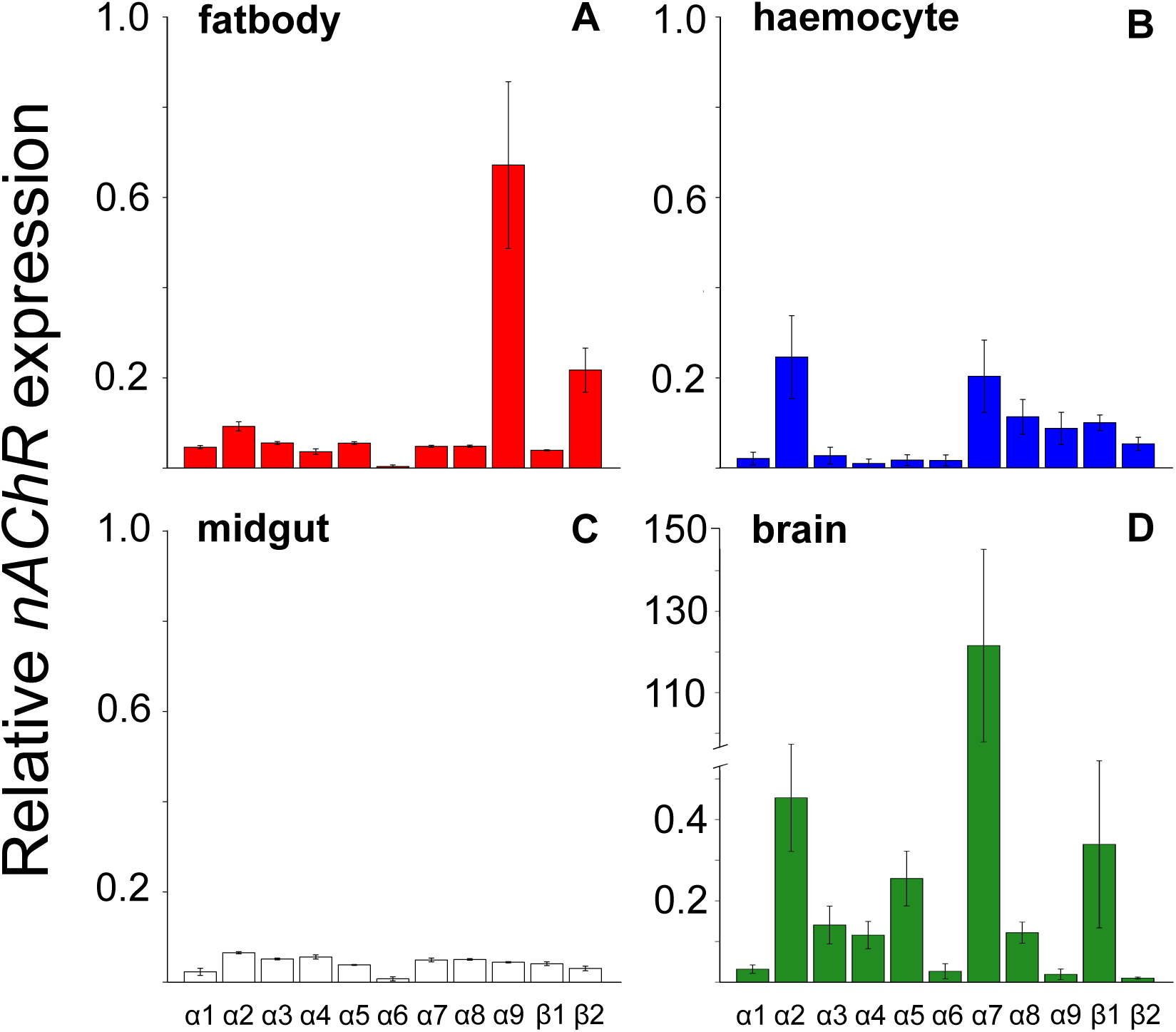
Relative expression of the nAChR subunits *α1-9, β1-2* normalized against rp49 in the four investigated tissue types: fatbody (A in red, N=7); haemocytes (B in blue, N=8); midgut (C in white, N=7); brain (D in green, N=5). There is an overall difference in the subunit expression between tissues (PERMANOVA: Pseudo-F_3,23_ = 7.76 P < 0.001). Error bars indicate SE.

**Figure 2:**
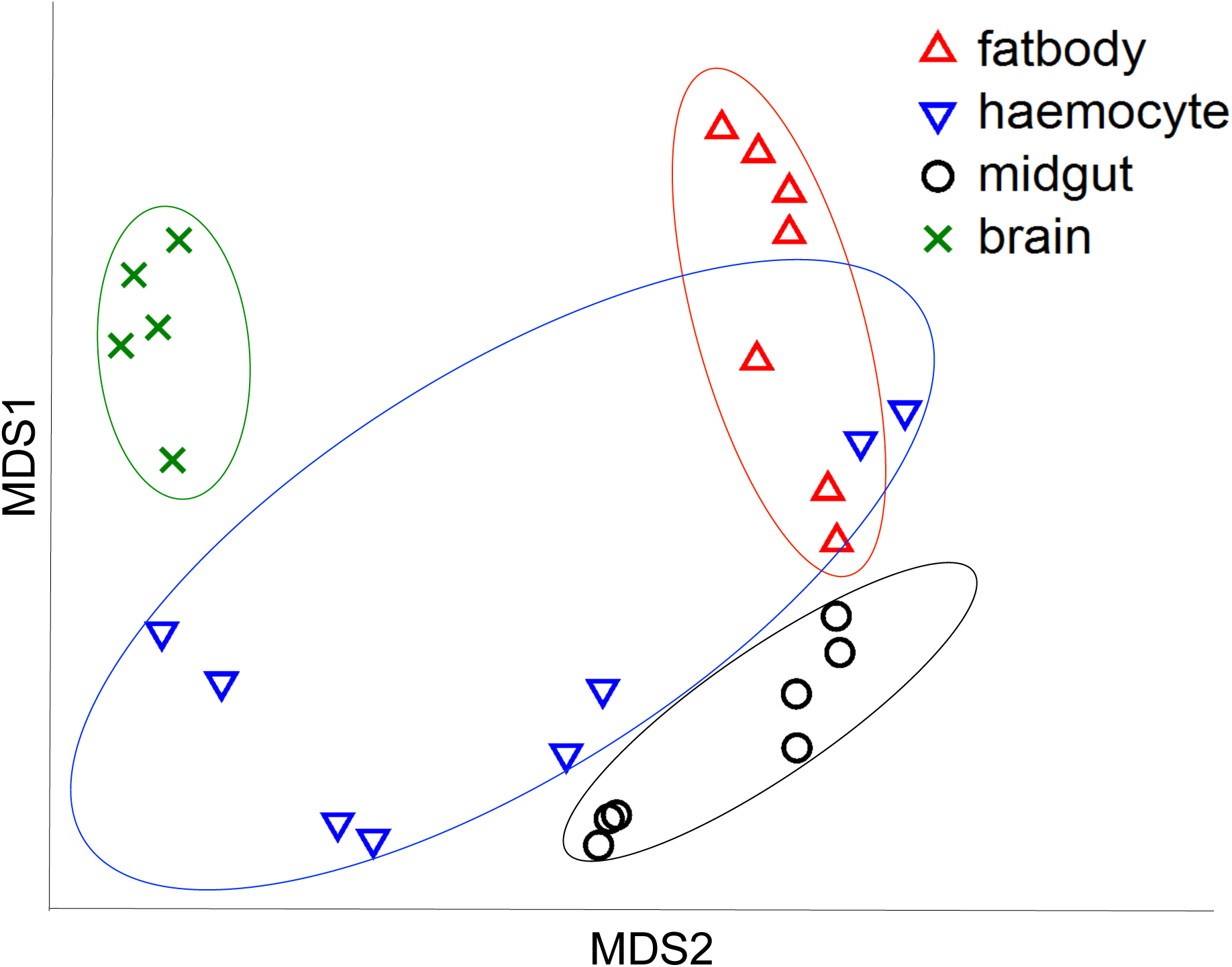
Multidimensional scaling (MDS) plot based on Chad distance of NAChR expression in four cell types (brain, haemocytes, fatbody and midgut) of honeybees. All groups varied significantly from each other (PERMANOVA pairwise comparison, all P < 0.001).

**Figure 3:**
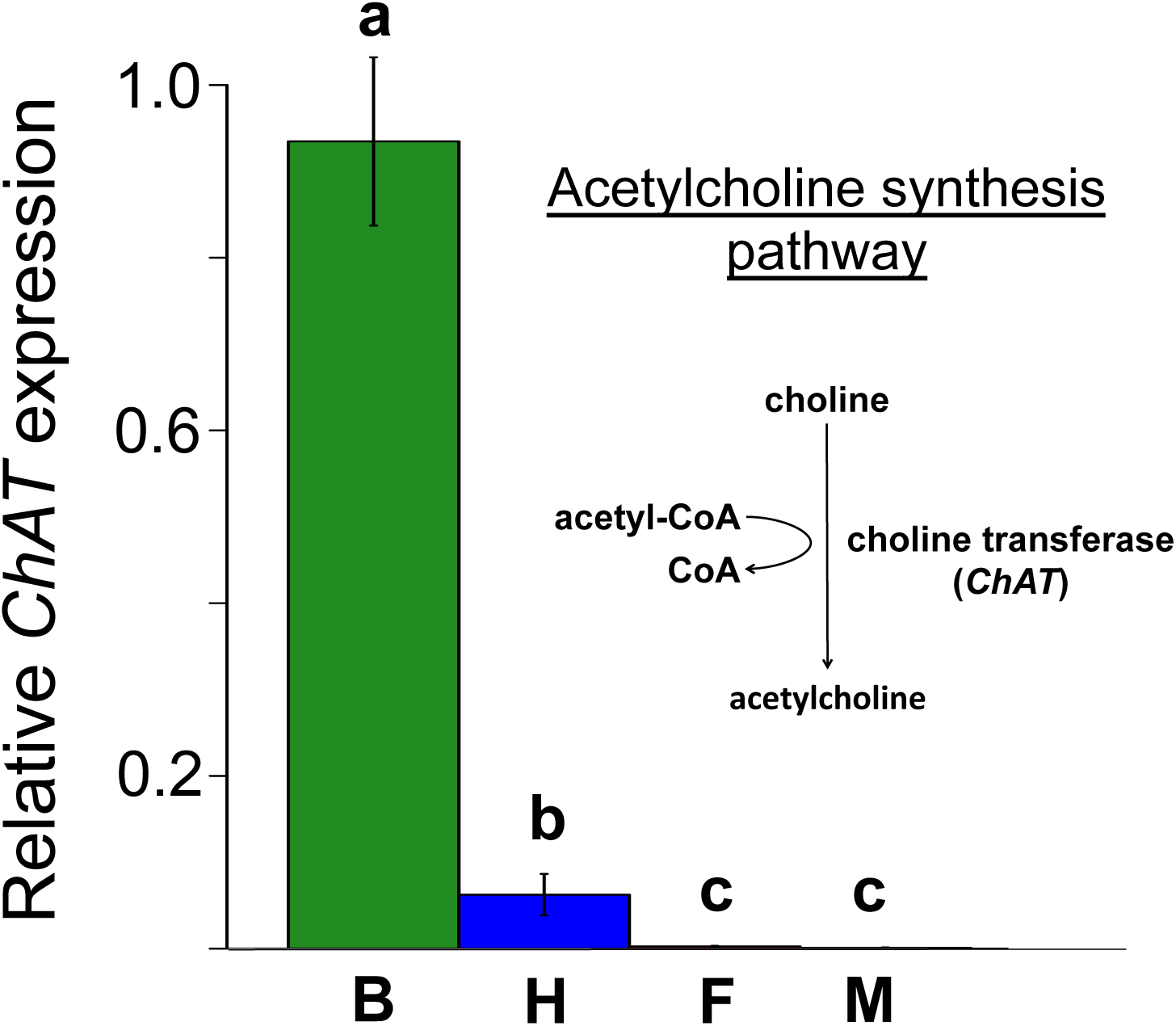
Mean ± s.e. relative expression of the *A. mellifera* choline transferase gene (*ChAT*) in four honeybee cell types: brain (green, B, N = 5); haemocytes (blue, H, N = 8); fatbody (red, F, N = 7); midgut (white, M, N = 7), and its role in acetylcholine synthesis. Only brain and haemocyte cells exhibit robust *ChAT* expression. Different letters indicate significant expression differences between cell types.

**Figure 4:**
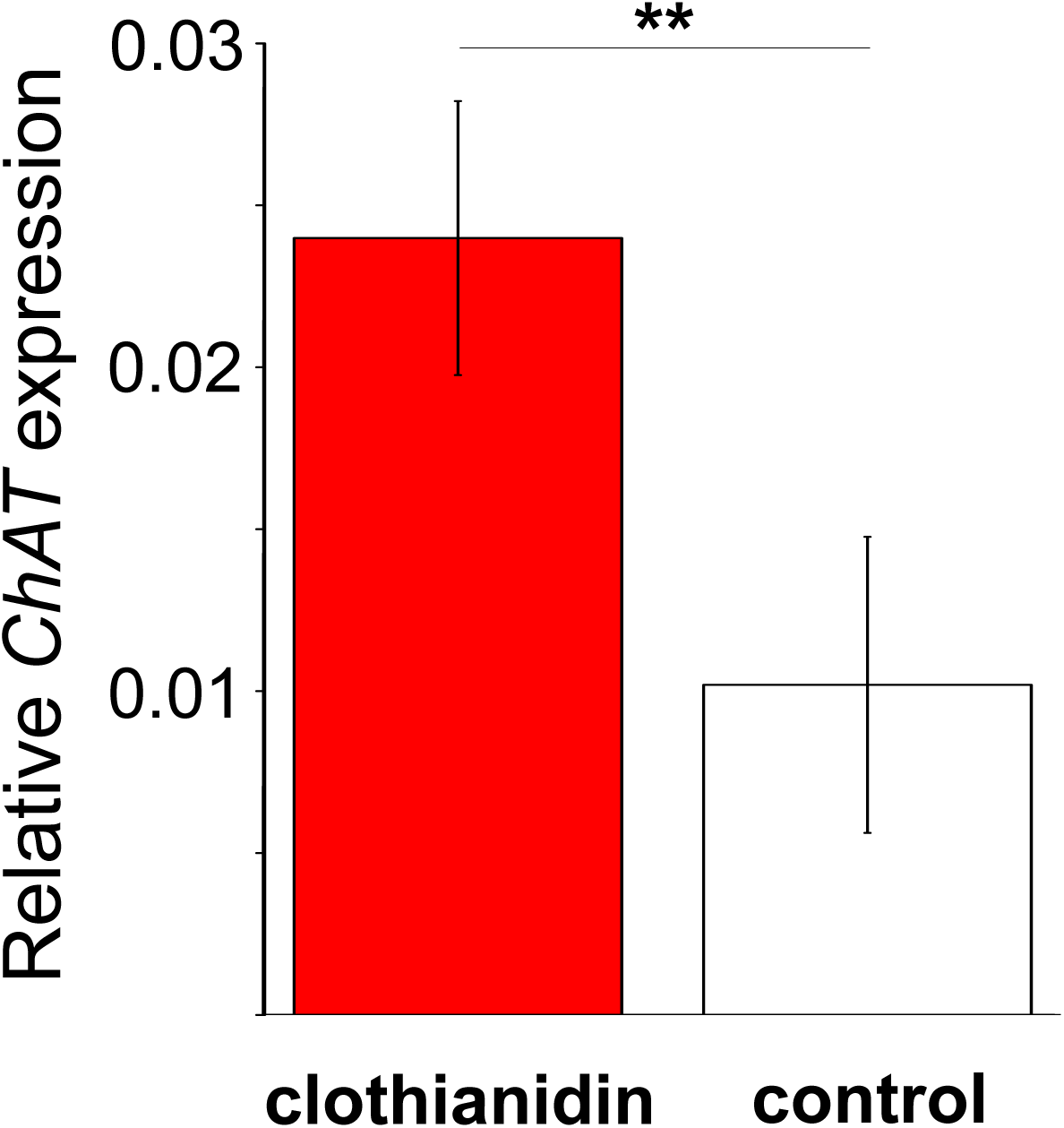
Mean ± s.e. relative expression of the choline transferase gene (*ChAT*) in the haemocytes of honeybees treated with neonicotinoid (clothianidin; red) or control (white).

## Discussion

In this study we demonstrate the widespread expression of *nAChR* subunits in non-neural and immune-relevant tissues in the honeybee *A. mellifera*. In addition we show that haemocytes in *A. mellifera* actively synthesize *ACh,* which strongly suggests *ACh*-based non-neural communication of the innate immune system in this important pollinator species. Lastly we experimentally establish that sub-lethal field relevant doses of the neonicotinoid clothianidin can influence this communication system *in vivo*.

Our results are in line with recent findings, which suggest the presence of non-neural and immune-related *ACh* based communication in a range of invertebrate model systems^18,21,23^. Work on pest insects indicates that, similar to our findings, different combinations of *nAChR* subunits are expressed in a wide range of non-neural tissues^22^. However, the expression of these sub-units by itself does not automatically indicate the presence of functional receptors, as demonstrated by Aztiria et al.^25^. Nevertheless, the fact that haemocytes can respond to the presence of *ACh* suggests that, at least in some species, functional receptors must be present^18,21^. In addition, haemocytes have been shown to synthesize acetylcholine-degrading enzymes (Acetylcholineesterase) likely terminating *ACh* based haemocyte excitation following pathogen exposure, thereby facilitating homeostasis of the immune system^19^. Our results strongly support these findings and indicate additionally that haemocytes are capable of actively synthesizing *ACh* themselves. Taken together these lines of evidence strongly suggest that invertebrate innate immune systems possess all essential components for sending, receiving and terminating *ACh* based signals. It is consequently likely that, similarly to their vertebrate counterparts^16^, the invertebrate innate immune system utilizes neural-independent *ACh-*based communication.

In addition, we show that secondary immune-relevant tissues, the fatbody and (to a lesser extent) the midgut, express *nAChR* subunits, which might be a common phenomenon in insects^22^. It is tempting to speculate that haemocytes could utilize *ACh* based signals to convey information to the fatbody and midgut and coordinate the systemic immune response during infections, orchestrating the cellular and humeral arm of the invertebrate innate immune system.

The utilization of *ACh*-based communication in the invertebrate immune system relies on the presence of functional *nAChR,* receptors that are the major target of neonicotinoid insecticides^26,27^. These neurotoxins exhibit high *nAChR* affinity in invertebrates, causing receptor overstimulation with lethal effects at even very low doses^24,28^. These systemic pesticides are of course aimed at pest species that eat plant tissue or suck sap, but, being systemic, they migrate into both pollen and nectar, so pollinators are exposed to them when visiting treated, flowering crops or contaminated wildflowers^29,30^. Once ingested, the pesticide is absorbed via the gut and passes through the haemolymph on the way to its designated target: the central nervous system^24^. In the haemolymph, neonicotinoids inevitably come into contact with haemocytes, with detrimental effects for haemocyte populations. It has been experimentally shown that neonicotinoids drastically decrease haemocyte numbers in the haemolymph of honeybees and also inhibit their ability to mount an effective immune response within 24 h of exposure^8^. In molluscs, the blocking of haemocyte-based *mAChR* before pathogen challenge promotes the expression of *Tumor Necrosis Factor* (TNF), which in turn results in elevated haemocyte apoptosis^21,31^. If a similar, *nAChR*-based, regulatory connection is present in the haemocytes of pollinators, *nAChR* blockage by neonicotinoids could directly explain their detrimental effects on haemocytes and by extension the immunosuppressive effects observed in honeybees^7^.

While the direct effects of neonicotinoids on neuronally associated traits like behaviour, memory and navigation^12,13,32^ are intuitively clear, the effects on other traits such as reproduction have not previously been adequately explained^33^. The finding that non-neural tissues can express *nAChR* could explain these counterintuitive effects by providing a mechanism for direct interaction with these tissues. It could also resolve a second puzzling phenomenon associated with neonicotinoid exposure: the susceptibility of insects to neonicotinoid exposure varies profoundly both within species (between different developmental stages)^34^, and between species^24^, as well as between studies using similar experimental set-ups^33,35^. In order to understand such effects, we have to consider the *ACh* (and by extension neonicotinoid) binding properties of *nAChRs*. These characteristics are determined by their subunit composition, which can greatly alter their binding probabilities and consequently also alter their susceptibility to neonicotinoid interference by orders of magnitude^24,36^. Work in vertebrates suggests that, in addition to species-specific and tissue-specific *nAChR* composition, environmental stimuli such as nicotine^37^ can alter receptor composition as well. If such results also hold for pollinators, then it may be that the species, tissue, time point and condition dependent *nAChR* composition might all influence the response to neonicotinoids, giving rise to the large variation in susceptibility observed both under natural and experimental conditions^22,33,35^.

In summary our results provide a novel, direct and mechanistically informed framework to understand the numerous unexplained and variable side effects associated with exposure of insects to sub-lethal doses of neurotoxic pesticides. In times of wild and managed pollinator decline such an analytical framework is urgently needed in order to identify, analyse and ultimately limit the side effects of pesticides.

## Methods

### Bee collection

Foraging *Apis mellifera* worker were collected between July and September 2016 on the campus of the University of Sussex, Brighton, UK (50°52’02.8"N 0°05’09.6"W). In all cases bees were collected between 09:00 and 11:00 in order to minimize gene expression variation caused by circadian rhythms. They were placed in 50 ml falcon tubes containing a moist cotton ball to provide them with water and to regulate relative humidity within the tube. The bees for Experiments 1 (tissue expression levels) were directly put on ice to cold anaesthetise them while the bees for Experiment 2 (clothianidin exposure) were placed in an dark incubator at 33°C and 80% relative humidity and provided with 60% sucrose water and kept at these conditions for 20 h for acclimatisation before the start of the experiment.

### Experiment 1: tissue-specific expression of nAChR subunits and ChAT

#### Tissue & RNA extraction

After cold immobilization (~10 min) the bees were decapitated using a sterile razor blade, dissected under RNA Later (Thermo Fisher) using a sterile dissection kit and either whole brain (N = 5), fatbody (N = 7) or midgut (N = 7) was extracted (one type of tissue per bee). For haemolymph extraction, the thorax and abdomen of the bees were carefully punctured after decapitation using a sterile dissection needle and haemolymph was collected using a sterile graded glass capillary. The haemolymph of two bees was pooled (total 16 bees; N = 8) and haemocytes were collected following standard protocol^38^. All tissues were homogenized in Trizol (ABI) using a sterile pestle and total RNA was extracted following the manufacturers instructions. The concentration and purity of RNA was determined on a Nanodrop 2000^®^.

### Experiment 2: clothianidin exposure

Sixty two foraging *A. mellifera* workers were randomly assigned to either treatment (N = 30) or control (N = 32). Following the 20 h acclimatisation period the feeders were removed. Four hours later the treatment group was provided with new feeders containing 60% sucrose solution spiked with 5 ppb clothianidin (using molecular grade acetone as solvent), while the control received sucrose solution with the same concentration of acetone only. All feeders were weighed before and after the experiment to the closest 0.001 g using a Kern PFB 300-3 scale to measure the dose (ng) of neonicotinoids that the bees had consumed. All bees had access to the feeders for 24 h after which haemolymph was collected from all surviving individuals, and samples of two bees were pooled (N = 7 treatment, N = 8 control) following the procedure of Experiment 1.

### RT transcription & qPCr analysis

100 ng of total RNA was used for reverse transcription using the Phuson RT-PCR kit (Thermo Scientific). Primers for all RT-qPCR assays of *α 1-9, β1-2, ChAT* and the reference gene *rp49* were designed using Primer3 ^39^ and published sequences available from GeneBank (See Tab.S3 for details). Primer efficiencies were measured using a dilution series of *Apis mellifera* brain cDNA (pooled subsamples of 5 individuals) covering three orders of magnitude including the cDNA concentration used in the reaction. Primer efficiencies were found to be above 91% for all primer pairs. Reaction specificity was confirmed by melting curve analysis. All analyses were performed on an ABI OneStep qPCR machine using SYBR green assays and were analysed using the OneStep software.

### Gene expression analysis

Gene expression analysis of the *nAChR* subunits *α1-9, β1-2, ChAT* and *rp49* were performed in 10 µl reactions using GoTaq^®^ qPCR Master Mix (Promega) and 0.5 µM of each specific primers (Sigma-Aldrich) on a StepOne™ Real-Time PCR Systems Applied Biosystems^®^ detection system. Samples of cDNA corresponding to 2 ng total RNA in 2 µL volumes were added and each sample analysed in technical duplicates. Each plate contained one negative control reaction for each primer pair using pooled and 1:10 diluted RNA extracts from 5 randomly chosen individuals in order to control for gDNA contamination. The following program was used for amplification: 95°C for 2 min, followed by 40 cycles of 30 s of 95°C denaturation, 30 s annealing at 59°C and 30 s extension at 72°C following by a melting curve to ensure PCR specificity. The data used for the analysis is the target gene expression normalized to the reference gene (*rp49*) expression.

### Data analysis

To compare the *nAChR* subunit expression patterns we used the programme PRIMER 6, version 6.1.13, + add-in, version 1.0.3 (PRIMER-E Ltd) to perform permutational multivariate analysis of variance (PERMANOVA) with the normalized relative expression of all 11 sub units as the response and tissue as the predictor variable. All tests were carried out using 9,999 permutations on a resemblance matrix using Chad distance estimates and the robustness of the results were tested using the Euclidian distance as an alternative estimate. We performed a SIMPER analysis to compare the expression of individual *nAChR* subunits according to tissue identity and tissue differentiation. All other tests were performed in R 3.2.4^40^. Survival was analysed as the proportion of bees that died over the duration of the experiment using a GLM with binomial data distribution. The other results of Experiment 2 were analysed using non-parametric statistics (Kruskal-Wallis and Wilcoxon tests) and Bonferroni corrections in cases of multiple testing. The MDS plot was generated in PRIMER 6; all other graphs were done in R using the sciplot package^41^.

